# Activity of the manganese efflux transporter SLC30A10 in dopaminergic but not GABAergic neurons protects against neurotoxicity

**DOI:** 10.1101/2022.02.21.481385

**Authors:** Cherish A. Taylor, Stephanie Grant, Thomas Jursa, Michael Aschner, Donald R. Smith, Reuben Gonzales, Somshuvra Mukhopadhyay

## Abstract

Metals such as copper, iron, and manganese (Mn) are essential for life, but induce neurotoxicity at elevated levels. Yet, the neuronal mechanisms of metal-induced neurological disease are largely unclear. A primary limitation has been an inability to selectively alter metal levels in specific neurons, so that the role of the targeted neurons in disease biology can be isolated. Here, we show that neuron-specific depletion of metal efflux transporters provides the feasibility to overcome this limitation by focusing on Mn, which accumulates in the basal ganglia and induces motor disease, and the Mn-specific efflux transporter SLC30A10. Pan-neuronal/glial *Slc30a10* knockout mice exhibited increased basal ganglia Mn levels and hypolocomotor deficits in early-life (pre-adulthood), which were exacerbated by Mn exposure. The locomotor deficits of the pan-neuronal/glial strain was associated with a reduction in evoked striatal dopamine release without dopaminergic (DAergic) neurodegeneration or changes in striatal tissue dopamine levels. Furthermore, DAergic-specific, but not GABAergic-specific, *Slc30a10* knockout mice recapitulated the hypolocomotor phenotype of the pan-neuronal/glial knockouts in early-life although Mn levels were elevated in the targeted basal ganglia regions of both the neuron-specific strains. Put together, our results imply that (1) activity of SLC30A10 in DAergic neurons is necessary to protect against early-life Mn neurotoxicity; (2) increasing Mn levels in DAergic neurons is sufficient to induce early-life motor disease, suggesting that Mn targets DAergic neurons in the early-life period to induce motor deficits; and (3) neuron-specific knockout of metal efflux transporters may be a widely applicable strategy to elucidate mechanisms of metal-induced neurotoxicity.

## Introduction

Essential metals such as copper (Cu), iron (Fe), and manganese (Mn) are required for normal brain development and function. However, elevated levels of these metals induce neurological disease ^1^. As examples, Wilson’s disease, a genetic disorder caused by mutations in the copper transporter *ATP7B*, is associated with the accumulation of copper in the brain and numerous neurological and psychiatric changes ^2^. Excess Fe accumulation in the substantia nigra is considered to be a risk factor for Parkinson’s disease ^3^. Elevated brain Mn levels induce a parkinsonian-like disorder in adults ^4-8^, and motor and executive function deficits in infants, children, and adolescents ^9-21^ (see below). Despite the clear relevance to human health, the specific neurons (or non-neuronal cell types) in the brain that are targeted by essential metals at elevated levels, and metal-induced changes in these cells that subsequently lead to neurological disease, remain largely unclear. A major limitation has been an inability to alter metal levels in a cell-type specific manner in the brain so that the role of the targeted cells in metal-induced disease can be determined. Notably, cellular levels of essential metals are tightly regulated by influx and efflux transporters, and metal efflux transporters exhibit a high degree of substrate specificity (as efflux is an energy-dependent process). Therefore, we hypothesized that cell-type specific depletion of metal efflux transporters in the central nervous system of human relevant animal models would provide a means to elucidate the neuronal targets and mechanisms of essential metal-induced neurological disease.

To test our hypothesis, we focus on Mn because of the occupational and environmental relevance of Mn neurotoxicity. Briefly, Mn neurotoxicity from occupational over-exposure in adults is a well-established cause of motor disease, which manifests as a parkinsonian-like disorder ^4-8^. Mn-induced parkinsonism is also recognized in individuals with compromised hepatic excretory function (e.g. due to cirrhosis), even in the absence of elevated Mn exposure, because Mn is primarily excreted by the liver ^22^. More recently, epidemiological studies have suggested that environmental Mn exposure (e.g. from sources such as drinking water or air) is an important public health problem, especially in the early-life periods of infancy, childhood and adolescence ^9-21^. In these developmentally-sensitive life stages, environmental Mn exposure is associated with substantial impairments of motor function, such as deficits in motor coordination, skilled motor function, tremor intensity, postural stability etc., and additionally, also induces executive function deficits ^9-21^. Currently, there are no treatments for Mn neurotoxicity, either in adults or early-life, and an understanding of the underlying neuronal mechanisms is essential for the development of effective interventions.

Prior work has established that excess Mn primarily accumulates in the basal ganglia to induce motor disease ^4-7^. In the basal ganglia, a network of predominantly dopaminergic (DAergic) or GABAergic neurons control movement, and degeneration or dysfunction of these neurons induce motor disease (e.g. Parkinson’s disease, due to degeneration of DAergic neurons in the substantia nigra pars compacta) ^23,24^. However, the fundamental question of whether Mn-induced motor disease reflects degeneration or dysfunction of basal ganglia DAergic or GABAergic neurons is unclear. Extensive mechanistic studies have focused on the parkinsonian phenotype of adults occupationally-exposed to Mn, and depending on experimental model or exposure regimen used, implicated divergent etiologies as the cause of the disease, including: (1) degeneration or dysfunction of GABAergic neurons; (2) degeneration of DAergic neurons; or (3) DAergic neuronal dysfunction without degeneration (reviewed in ^5,6,8^). Other mechanisms that would be expected to impact basal ganglia DAergic or GABAergic neurons to produce motor deficits (e.g. glutamatergic excitotoxicity, cholinergic dysfunction or activation of glial cells) have also been implicated ^5,6,8^. In contrast with the considerable, albeit conflicting, literature about Mn-induced parkinsonism in adults, there is little to no understanding of how early-life Mn exposure induces motor deficits. This is primarily because the focus of the few mechanistic studies on environmental Mn neurotoxicity has been on the executive function deficits that usually co-occur with motor dysfunction ^17^. While some of these studies attributed the executive domain deficits to Mn effects on the prefrontal cortex ^25,26^, insights about motor deficits did not emerge. Overall, the as yet unclear effects of elevated Mn levels on basal ganglia DAergic or GABAergic neurons that lead to motor disease represent a major limitation in our understanding of the neurotoxic effects of Mn.

To discriminate between the contribution of DAergic and GABAergic neurons to Mn-induced motor disease, in this study, we compared the neuromotor and neurotoxic consequences of elevating Mn levels only in DAergic or GABAergic neurons with Mn level elevations in all neurons using genetically-modified mice. Our mouse models are based on the neuron-specific or pan-neuronal/glial knockout of the critical Mn efflux transporter, SLC30A10 ^27-32^. Briefly, in 2012, mutations in *SLC30A10* were reported to induce the first hereditary form of Mn-induced neurotoxicity in humans ^33-35^. Over the last few years, our work characterized SLC30A10 to be a specific Mn efflux transporter that reduces cellular Mn levels to protect against Mn toxicity ^28,29,31,32,36^. Our studies in whole body and tissue-specific *Slc30a10* knockout mice revealed that an important function of SLC30A10 at the organism level is to directly mediate hepatic and intestinal Mn excretion, which prevents accumulation of Mn in the body, including in the brain ^29,30^. Interestingly, we had also detected SLC30A10 expression in the basal ganglia, raising the possibility that, in addition to its role in Mn excretion, SLC30A10 has a second function in directly controlling basal ganglia Mn levels that may be leveraged to elucidate the neuronal targets and mechanisms of Mn-induced motor disease. Here, we show that (1) pan-neuronal/glial *Slc30a10* knockouts exhibit elevated basal ganglia Mn levels and develop motor deficits in early-life, which are enhanced by elevated Mn exposure; (2) the motor deficits in the pan-neuronal/glial strain is associated with a reduction in striatal evoked dopamine release, but there is no evidence of DAergic neurodegeneration; and (3) DAergic-specific, but not GABAergic-specific *Slc30a10* knockout mice phenocopy the pan-neuronal/glial strain and exhibit early-life motor deficits, even though Mn levels are elevated in targeted basal ganglia neurons/regions of both the neuron-specific strains. The totality of our findings (1) characterize the activity of SLC30A10 in DAergic neurons as vital for neuroprotection; (2) identify DAergic neurons to be a primary target of early-life Mn toxicity; and (3) suggest that neuron-specific knockout of metal efflux transporters in mice may be broadly applicable to study the mechanisms of metal-induced neurological disease.

## Results

### Activity of SLC30A10 in the basal ganglia is necessary to reduce Mn levels in early-life

We previously reported that SLC30A10 is widely expressed in the brain, including in the basal ganglia (striatum, globus pallidus, and substantia nigra) and thalamus ^30^. RNAscope imaging confirmed SLC30A10 expression in DAergic neurons of the substantia nigra pars compacta and GABAergic neurons of the striatum (not shown). To determine whether expression of SLC30A10 in the brain/basal ganglia has a direct role in regulating Mn homeostasis and neurotoxicity, we used pan-neuronal/glial *Slc30a10* knockout mice (*Slc30a10*^*fl/fl*^;Nes-*Cre*) that we have previously reported ^30^. This brain tissue-specific knockout was generated by crossing floxed *Slc30a10* mice with Nestin-*Cre*, which induces recombination in all neurons as well as glial precursors ^30^. We reconfirmed specificity of the knockout in this study (**Fig.1A**; we previously demonstrated that *Slc30a10* is depleted in the basal ganglia and thalamus of the pan-neuronal/glial knockout strain ^30^). As Nestin-*Cre* is active during embryonic development, we anticipated that the pan-neuronal/glial *Slc30a10* knockout mice would be ideal for elucidating the role of brain SLC30A10 expression in the homeostatic control and toxicity of Mn in the early-life period (defined as PND 1-60 in mice, which corresponds to human infancy, childhood and adolescence ^37,38^; mice reach adulthood at PND 60 ^37,38^). Metal analyses using brain punch tissue samples and inductively coupled plasma-mass spectrometry (ICP-MS) at PND 21 revealed that, in comparison with littermate controls and in the absence of Mn exposure, Mn levels in the basal ganglia (striatum, globus pallidus, and substantia nigra) and thalamus of the pan-neuronal/glial knockouts had a non-significant trend towards an increase (**Fig.1B**). Low-dose daily oral Mn treatment from birth, modeling human environmental Mn exposure, produced ∼3-6 fold increase in basal ganglia and thalamus Mn levels (**Fig.1B**). Under these exposure conditions, Mn levels of the pan-neuronal knockouts were significantly higher than littermates (by ∼75%) (**Fig.1B**). Note that, in addition to the basal ganglia regions, we included the thalamus in these assays because it expresses SLC30A10 ^30^ and receives the basal ganglia output ^23,24^, and we conducted the experiment at PND 21 because previous studies show that the effect of oral Mn exposure on brain Mn levels is highest in early postnatal life (i.e. around PND 21) before Mn excretory capacity of rodents is fully developed ^22,39-41^. Importantly, Mn treatment also increased blood Mn levels at PND 21, but there was no difference between pan-neuronal/glial knockouts and littermate controls (**Fig.1C**), implying that Mn level increases observed in the basal ganglia and thalamus of the pan-neuronal/glial knockouts were independent of the excretory function of SLC30A10. Additional analyses in 3 month old pan-neuronal/glial knockout and littermate control mice that received daily oral Mn from birth revealed a much smaller Mn treatment effect on brain or blood Mn levels (≤1-2 fold; compared to ∼3-6 fold at PND 21) and no genotype-specific differences (**Fig.1D&E**), despite the longer period of exposure. Results from these adult animals are consistent with prior work on the age-dependent relationship of oral Mn exposure and increased body Mn burden ^41^ and suggest that phenotypic changes observed in the pan-neuronal/glial *Slc30a10* knockouts reflect Mn level changes in the early postnatal period. Levels of other metals (Fe, Cu, and Zn) were essentially similar across genotypes at both PND 21 and 3 months (**Fig.S1A&B**), validating the specificity of the changes in Mn. Overall, the metal analyses data indicate that activity of SLC30A10 in the basal ganglia and thalamus is required to regulate Mn levels in these brain regions in early-life.

**Figure 1.**
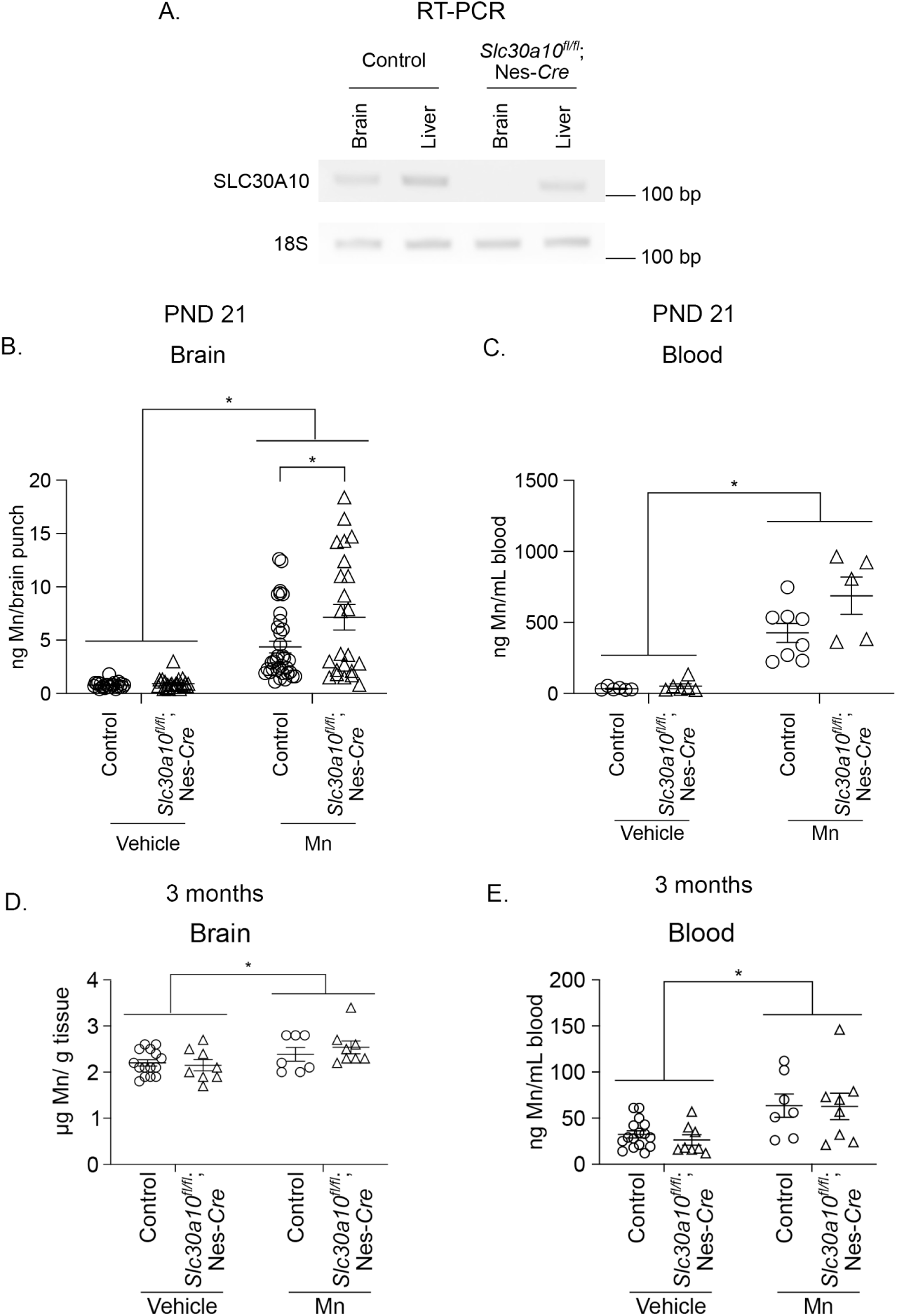
Mn levels in the basal ganglia and thalamus are elevated in pan-neuronal/glial *Slc30a10* knockout mice (*Slc30a10*^*fl/fl*^;Nes-*Cre*) in early postnatal life. ***A***. RT-PCR to detect SLC30A10 or 18S mRNA from brain (midbrain containing the basal ganglia) and liver tissue of pan-neuronal/glial *Slc30a10* knockouts or littermate controls. ***B-E***. ICP-MS to measure Mn levels in brain punches from the striatum, globus pallidus, substantia nigra, and thalamus (***B***), blood (***C*** and ***E***), or midbrain containing the basal ganglia (***D***) in pan-neuronal/glial *Slc30a10* knockout or littermate control mice treated with daily vehicle or Mn (10 mg Mn/kg body weight per day) and euthanized at PND 21 (***B*** and ***C***) or 3 months (***D*** and ***E***). For ***B***, n=23-35 samples for each genotype and Mn-treatment condition (group) obtained from 5-8 mice per group (each brain region was sampled once per animal). For ***C***, n=5-8 mice per group. For ***D*** and ***E***, n=7-16 mice per group. Mean ± SE. *, P < 0.05 using two-way ANOVA and Tukey’s *post hoc* test.

### Activity of SLC30A10 in the brain protects against Mn-induced motor deficits in early-life

We then sought to determine whether the Mn level elevations in the basal ganglia and thalamus of the pan-neuronal/glial knockouts produced motor deficits using the open-field test. These assays were performed in adolescent mice at 7-8 weeks of age with or without Mn exposure. There were no differences between genotype and treatment conditions in the first 5 min of the test (**Fig.2**), which reflects reactivity and exploratory behavior of an animal in a novel environment. Notably, even in the absence of Mn exposure and compared with littermate controls, pan-neuronal/glial knockouts exhibited reduced movement in the next 10 min interval (**Fig.2**), which measures generalized locomotor activity. As expected, Mn treatment had an inhibitory effect on activity in the second 10 min interval irrespective of genotype (**Fig.2**). Importantly, after Mn treatment, the locomotor activity of pan-neuronal/glial knockouts was still less than that of littermate controls (**Fig.2**). Results in **Figs.1&2**, put together, indicate that SLC30A10 is active in the basal ganglia, where it regulates Mn levels to protect against Mn-induced motor deficits in early-life.

**Figure 2.**
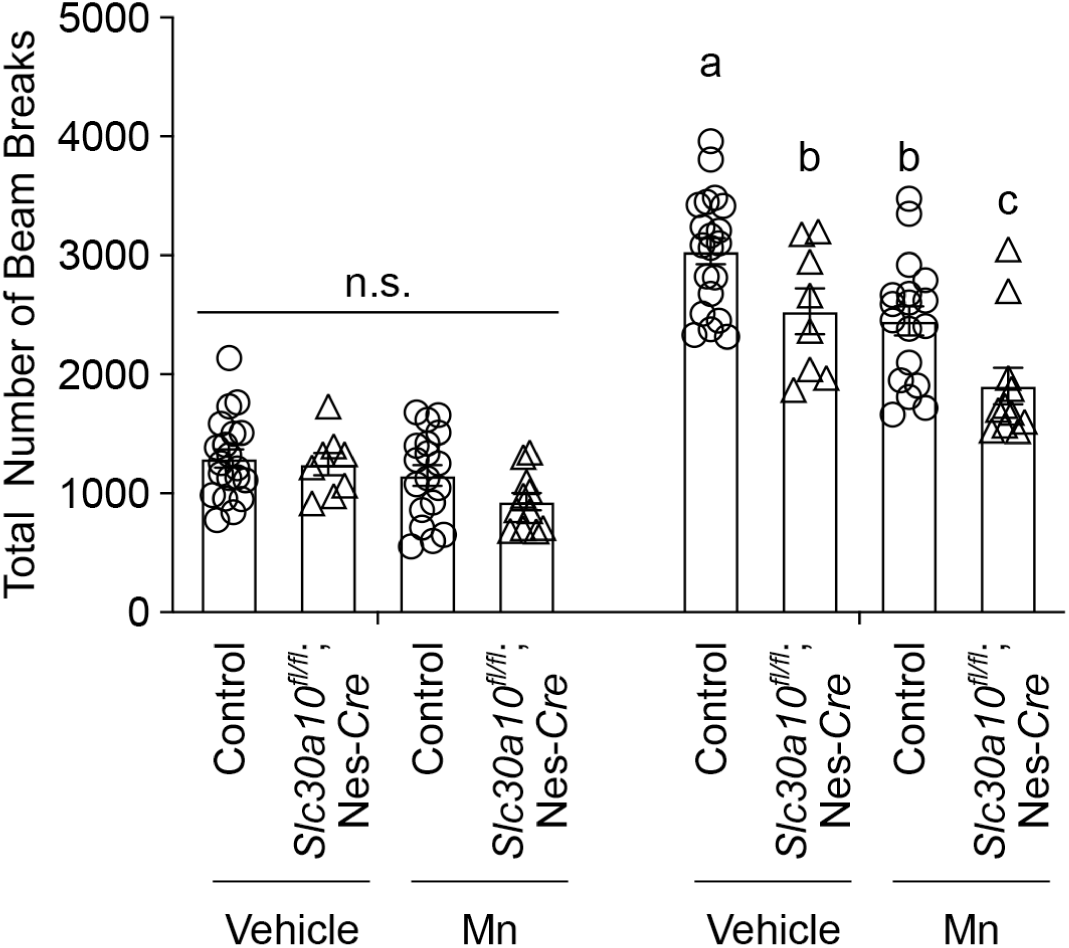
Pan-neuronal/glial *Slc30a10* knockout mice (*Slc30a10*^*fl/fl*^;Nes-*Cre*) develop motor deficits in early life. The open field test was used to monitor activity of 7-8 week old pan-neuronal/glial knockouts or littermate controls with daily vehicle or Mn treatment from birth. N = 20, 8, 18, and 11 for vehicle- or Mn-treated controls or knockouts, respectively. Mean ± SE. Unique letters notate p < 0.05 using repeated measures two-way ANOVA and Tukey’s *post hoc* test. n.s., not significant.

### Pan-neuronal/glial Slc30a10 knockouts exhibit a deficit in evoked dopamine release without DAergic neurodegeneration

To gain insights into the mechanisms that underlie the hypolocomotor phenotype of the pan-neuronal/glial *Slc30a10* knockouts, we performed *in vivo* microdialysis experiments to assay for functional changes in neurotransmitter release; tissue neurochemistry experiments to quantify total neurotransmitter levels; and microscopy analyses to determine whether neurodegenerative changes were evident. We focused on the DAergic system given the likelihood that a decrease in dopamine release or production in the basal ganglia would produce locomotor deficits. Furthermore, we performed these experiments without Mn exposure because the pan-neuronal/glial knockouts exhibited a hypolocomotor phenotype in the absence of Mn treatment (**Fig.2**). Finally, for logistical reasons, these assays were performed in adult mice, but as described above, phenotypic changes in the pan-neuronal/glial knockouts at this age are reflective of Mn level increases in the basal ganglia in the early post-natal period as brain Mn levels at 3 months are comparable between the knockouts and littermate controls (**Fig.1D**).

*In vivo* microdialysis assays revealed that, compared with littermate controls, evoked release of dopamine in the striatum was markedly reduced in the pan-neuronal/glial knockouts (**Fig.3A&C**). Baseline dopamine levels were, however, comparable between the genotypes (**Fig.3A&B**). Additionally, there were no genotype-specific changes in the total levels of striatal dopamine or the dopamine metabolite DOPAC (**Fig.3D**) or evidence of DAergic neurodegeneration in the pan-neuronal/glial knockouts (**Fig.3E**). The totality of the neurochemical and microscopy results indicate that a deficit in potassium-evoked dopamine release, but not DAergic neurodegeneration or changes in bulk tissue dopamine content, leads to the hypolocomotor phenotype of the pan-neuronal/glial *Slc30a10* knockout mice.

**Figure 3.**
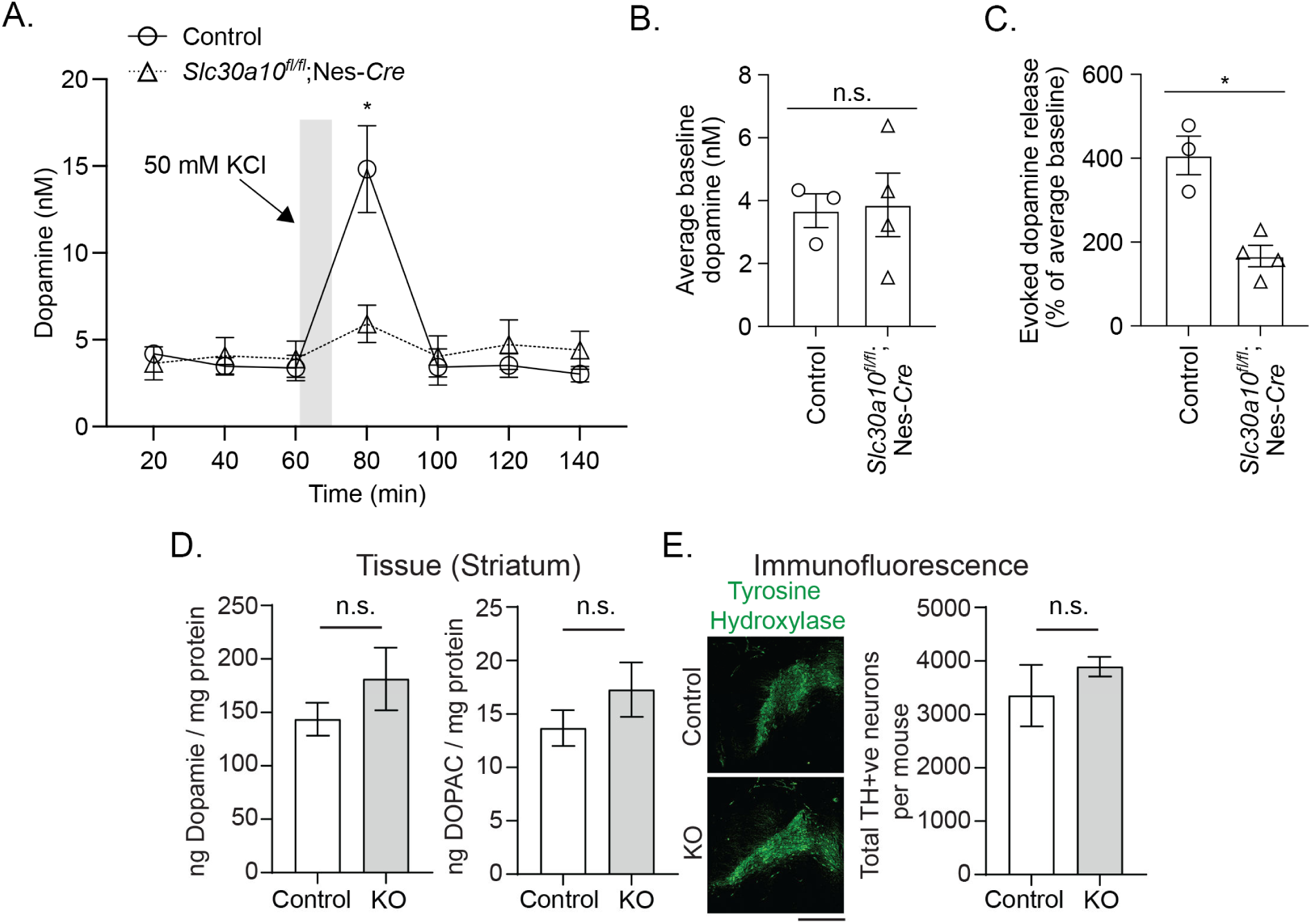
Evoked dopamine release is inhibited in pan-neuronal/glial *Slc30a10* knockout mice (*Slc30a10*^*fl/fl*^;Nes-*Cre* or KO). ***A***. *In vivo* microdialysis to measure dopamine levels in the dorsal striatum of 3-4 month old vehicle-treated pan-neuronal/glial knockouts or littermate controls. N=3-4 mice per genotype. Mean ± SE. *, p < 0.05 using repeated measures two-way ANOVA and Sidak’s *post hoc* test for the difference between genotypes at the 80 min time-point. ***B***. The average of the first three baseline dopamine measures from ***A***. Mean ± SE. n.s., not significant by *t* test. ***C***. The percent increase in KCl-evoked dopamine release from ***A***. Mean ± SE. *, p < 0.05 by *t* test. ***D***. Striatal tissue dopamine or DOPAC levels in vehicle-treated 3-month old pan-neuronal/glial knockouts (KO) or littermate controls. N=7-8 mice per genotype. Mean ± SE. n.s., not significant by *t* test. ***E***. Immunofluorescence to detect tyrosine hydroxylase (TH)-positive neurons in vehicle-treated pan-neuronal/glial knockout mice (KO) or littermate controls with quantification from N=3 mice per group. Mean ± SE. n.s., not significant by *t* test. Bar, 500 µm.

### Mn levels are elevated in targeted regions/neurons of dopaminergic- or GABAergic-specific Slc30a10 knockout mice

A reduction in dopamine release may occur due to a primary injury in DAergic neurons or changes in GABAergic neurotransmission ^42^. Thus, the neurochemistry results could not isolate which neuronal subtype was primarily driving the hypolocomotor phenotype of the pan-neuronal/glial knockouts. To differentiate between the roles of DAergic and GABAergic neurons, we generated DAergic- or GABAergic-specific *Slc30a10* knockout mice by crossing floxed *Slc30a10* mice with TH- or GAD2-*Cre* ^43-45^. To validate specificity of knockout, we performed qRT-PCR assays using brain punch samples. In the DAergic-specific strain, SLC30A10 was depleted in the substantia nigra, which is enriched in DAergic neurons, but not the striatum, which contains GABAergic neurons, while the opposite effect was observed in the GABAergic strain (**Fig.4A**). A more extensive depletion of SLC30A10 in the substantia nigra of the DAergic strain was not observed (**Fig.4A**) likely because bulk brain tissue punches from the substantia nigra include the pars compacta, which contains targeted DAergic neurons, and the pars reticulata, which is enriched in non-targeted GABAergic neurons. These expression studies suggested that our conventional approach of performing ICP-MS Mn analyses on brain punch samples would not have the spatial resolution necessary to detect neuron/region-specific changes in Mn levels in these neuron-specific knockout strains. Therefore, we performed laser ablation-ICP-MS (LA-ICP-MS), which provides a measurement spatial resolution of ∼20 µm, on brain sections to spatially resolve Mn levels in targeted regions/neurons. Analyses at PND 21 confirmed that Mn levels were elevated only in targeted regions of the neuron-specific strains (**Fig.4B**). Compared with littermate controls, the DAergic-specific knockouts exhibited increased Mn levels in the substantia nigra pars compacta (predominantly DAergic) but not surrounding regions including the substantia nigra par reticulata (predominantly GABAergic) and the retromammillary nucleus (**Fig.4B**). Similarly, the GABAergic-specific knockouts exhibited increased Mn levels in the striatum and globus pallidus (predominantly GABAergic), but not in the cortex (**Fig.4B**). Thus, neuron-specific *Slc30a10* knockout mice exhibit region/neuron-specific loss of SLC30A10 and elevation of Mn levels.

**Figure 4.**
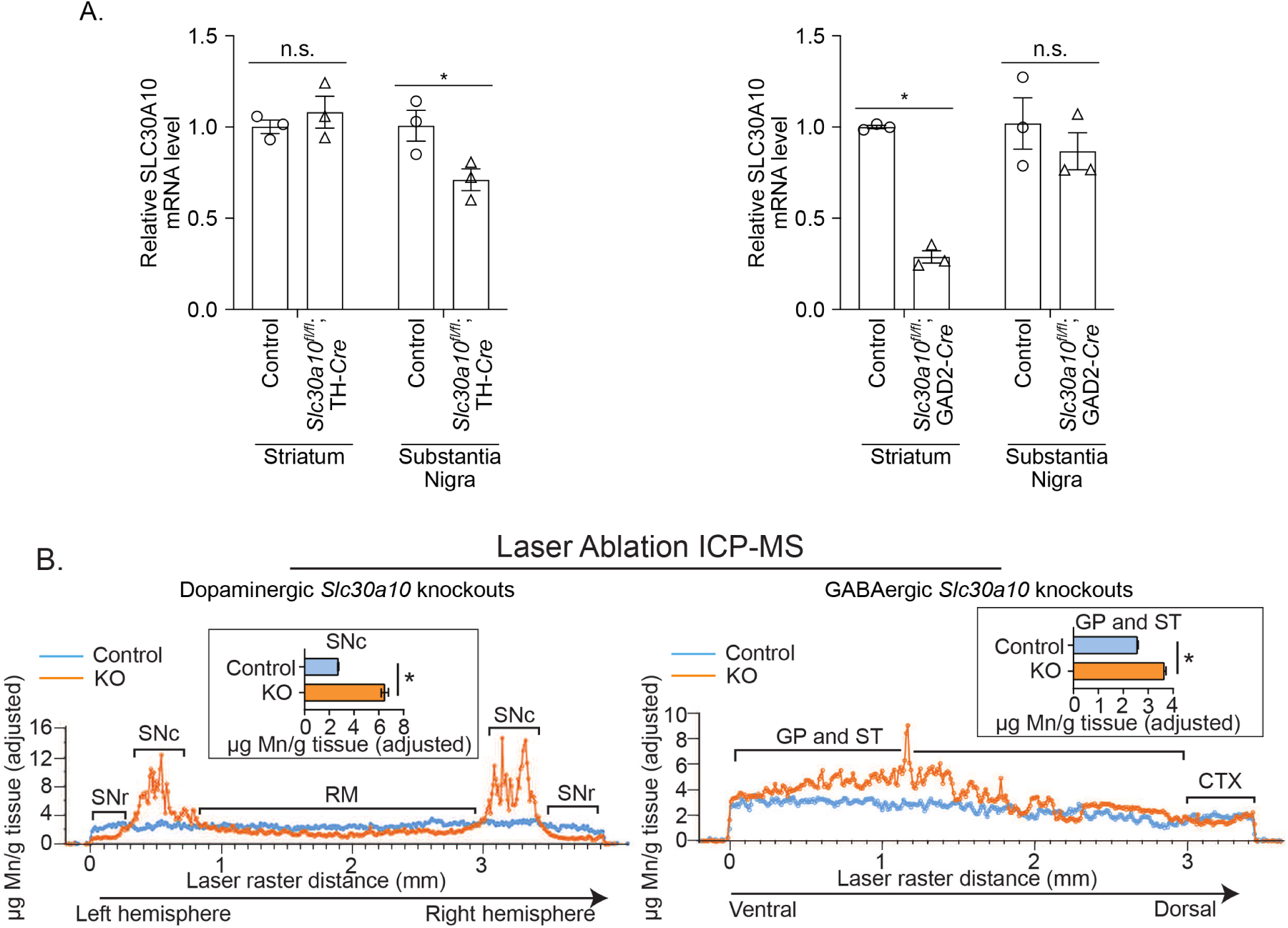
SLC30A10 is depleted and Mn levels are elevated in targeted regions of DAergic- or GABAergic-specific *Slc30a10* knockout mice. ***A***. QRT-PCR from brain punches of DAergic-specific or GABAergic-specific *Slc30a10* knockout mice (*Slc30a10*^*fl/fl*^;TH-*Cre* or *Slc30a10*^*fl/fl*^;GAD2-*Cre*, respectively) and their littermate controls. Mean expression in striatum and substantia nigra of each littermate control strain is independently normalized to 1. N=3 mice per condition. Mean ± SE. *P<0.05 by two-way ANOVA and Sidak’s *post hoc* test. ***B***. LA-ICP-MS chromatograms and quantification from PND 21 mice (littermate control and DAergic-specific or GABAergic-specific *Slc30a10* knockouts (KO)) treated with daily oral Mn (10 mg/kg) from PND 1. Analysis used a 20 µm laser spot size rastered across 20 µm thick brain slices. For the DAergic strain, rastering was across both hemispheres and traversed the substantia nigra pars reticulata (SNr), SN pars compacta (SNc), and retromammamillary nucleus (RM). For the GABAergic strain, rastering was from the ventral to dorsal aspect of one hemisphere and traversed the globus pallidus (GP), striatum (ST), and cortex (CTX). Regions were identified using an atlas and neuronal identity verified via RNAScope imaging of alternating sections. Inset bar graphs are mean (±SE) Mn levels for SNc or GP and ST calculated from the respective chromatograms (n=92 data points for SNc and 296 for GP and ST; * P<0.05 by *t*-test).

### Dopaminergic-, but not GABAergic-, specific Slc30a10 knockout mice develop early-life hypolocomotor deficits

We then assayed for the locomotor activity of the neuron-specific strains using the open-field test. Assays were performed in adolescent mice at 7-8 weeks of age, which corresponds to the age at which pan-neuronal/glial *Slc30a10* knockouts were analyzed, and without Mn exposure, because the pan-neuronal/glial knockouts exhibited locomotor deficits in the absence of Mn treatment (**Fig.2**). In the first 5 min interval, activity of both the neuron-specific knockouts were comparable to respective littermate controls (**Fig.5A&B**). Notably, in the second 10 min interval, the DAergic-specific knockout exhibited hypolocomotor deficits, similar to the pan-neuronal/glial strain (**Fig.5A**), but activity of the GABAergic-specific knockout was not affected (**Fig.5B**). Thus, loss of SLC30A10 and increase in Mn levels in DAergic, but not GABAergic, neurons produces motor deficits in early-life, suggesting that Mn targets DAergic neurons to induce motor disease in the early-life period.

**Figure 5.**
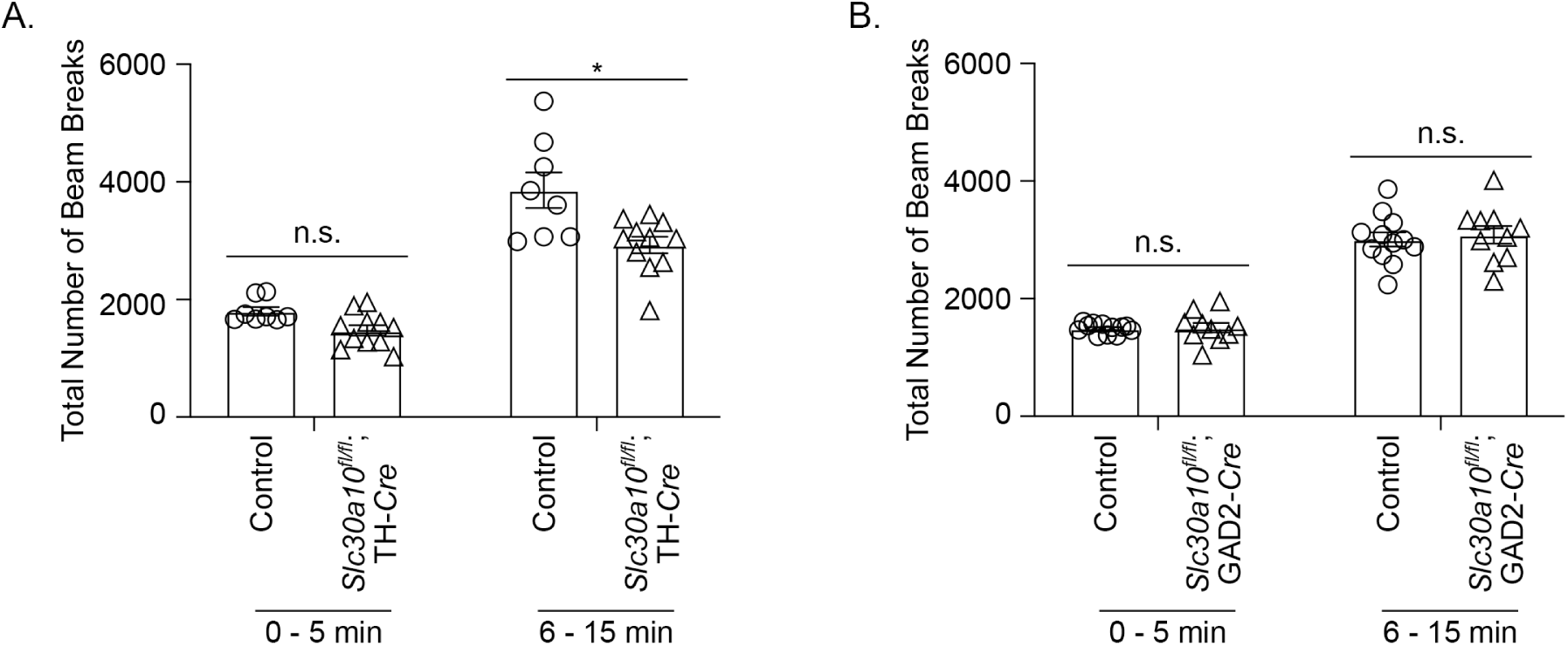
DAergic-specific, but not GABAergic-specific, *Slc30a10* knockout mice develop early-life motor deficits. ***A*** and ***B***. The open field test was used to monitor activity of 7-8 week old DAergic-specific (***A***) or GABAergic-specific (***B***) *Slc30a10* knockout mice (*Slc30a10*^*fl/fl*^;TH-*Cre* or (*Slc30a10*^*fl/fl*^;GAD2-*Cre*, respectively) and their littermate controls. In ***A***, n = 8 controls and 11 knockouts. In ***B***, n = 12 controls and 10 knockouts. Mean ± SE. *, p < 0.05 using repeated measures two-way ANOVA and Sidak’s *post hoc* test.

## Methods

### Generation of knockout strains, age of animals, and husbandry

All mouse experiments were approved by the Institutional Animal Care and Use Committee of the University of Texas at Austin (Austin, TX). We previously described the development and characterization of homozygous floxed *Slc30a10* mice (*Slc30a10*^*fl/fl*^), in which exon 1 of *Slc30a10* is flanked by loxP sites, and pan-neuronal/glial *Slc30a10* knockouts ^27,29,30^. DAergic- or GABAergic-specific *Slc30a10* knockouts were generated by crossing *Slc30a10*^*fl/fl*^ mice with TH-*Cre* (Jackson Laboratories strain number 008601) or GAD2-*Cre* (Jackson Laboratories strain number 019022), respectively, which are widely used to target DAergic or GABAergic neurons ^43-45^. Breeding, genotyping, and animal housing were as described previously ^27,29,30,46^.

Epidemiological studies indicate that the early life stages of infancy, childhood and adolescence in humans are most relevant for environmental Mn neurotoxicity ^9-21,37,38^. In mice, these stages correspond to PND 1-60, with PND 1-21 representing the early postnatal period, and PND 21-60 representing the prepubescent-adolescent period. We refer to this PND 1-60 age range as “early-life”, and performed metal analyses and behavioral assays within this range. Inclusion of the adolescent period is notable as it is often overlooked in mechanistic studies.

### RT-PCR, quantitative RT-PCR, and brain punching

These were performed as described previously^27,29,30,46^.

### Oral Mn treatment

Mn-treated mice daily received 50 mg/kg MnCl_2._4H_2_0 per day (∼10 mg absolute Mn/kg per day) beginning at birth and continuing until euthanasia. For pre-weaning exposure (birth – PND 21), Mn was administered by hand via pipette in a total volume of 5 µL (birth – PND 6) or 10 µL (PND 7 - 20). A 250 mg/mL MnCl_2_.4H_2_O stock was diluted in water containing 2.5% of the natural sweetener stevia, and the required dose, based on the pup’s weight, was delivered. Vehicle-treated animals received water containing 2.5% stevia. Pre-weaning Mn delivery did not impact nursing by the dam or milk intake by the pup. Post-weaning Mn exposure was administered via drinking water containing 0.2 mg Mn/ml, based on the fact that animals’ total water intake is ∼25% of body weight, and the above Mn level provides the required 10 mg Mn/kg daily dose. Animal body weights and water intake were measured regularly, and Mn concentration was adjusted if necessary to ensure target dose delivery. Vehicle-treated animals received only water in the post-weaning period.

We used the above oral Mn exposure regimen because drinking water-based oral exposure is a major source of environmental Mn exposure in human infants, children, and adolescents ^36^. The regimen used by us, and related similar regimens, (1) approximate the increase in Mn exposure in infants and children consuming well water contaminated with 1.5 mg Mn/L, which is the median well water concentration associated with neurological deficits in children ^36^; (2) induce modest increases in brain Mn that are comparable to humans ^36^; and (3) produce measurable behavioral deficits without overt toxicity ^36^.

### Open Field Test

The open field test was performed exactly as described by us previously ^30,46^. Animals were placed individually in the open field chamber (Opto-Varimex 4 Activity Meter, Columbus Instruments, Columbus, OH) for 15 min. Data was binned into two intervals: one 5-min interval and one 10-min interval. The first 5 min provides information about the animal’s exploratory behavior, while the second 10 min provides information about general locomotion. The test was performed during the light period of the animals’ light-dark cycle. The experimenter was not in the room during testing. Once testing was complete, animals were removed from the chamber and returned to their home cage. The open field chamber was cleaned with 5% ethanol between animals.

### Immunofluorescence Assays

Whole brain was collected and post-fixed in 4% paraformaldehyde for at least 48 hours. Following fixation, tissue was placed in a 30% sucrose solution until the brains sunk (∼48 hours), then placed in 1X phosphate-buffered saline (PBS). For sectioning, whole brains were embedded in Optimal Cutting Temperature (OCT) compound and frozen in cold isopentane (Thermo Fischer Scientific). Tissue was sliced on a cryostat (set to ∼-18°C) in 20 µm sections. Sections were placed free-floating in 1X PBS and stored at 4°C until staining. For staining, sections were transferred to a 24-well plate with one section per well and processed in the following order: three 10 min washes in 1X PBS at room temperature, 1 h incubation in 1% triton (in PBS) at room temperature, 1 h incubation in blocking solution (see below) at room temperature, overnight incubation in the primary antibody at 4°C, five 10 min 1X PBS washes at room temperature, 2 h incubation in the secondary antibody at room temperature, 10 min incubation in DAPI, and five 10-min 1X PBS washes at room temperature. Sections were placed on a shaker for each step. Blocking solution was made as previously described ^27,28,30^. Primary anti-TH antibody (Immunostar, Cat# 22941), secondary antibody, and DAPI were diluted 1:1000 in blocking solution. After staining, sections were placed on glass slides. Coverslips were placed over the sections using mounting media (90% glycerol in Tris pH 7.4) and sealed with clear nail polish. Images were captured using a Nikon swept-field confocal with an inverted Nikon TiE inverted microscope. The objective was a Nikon 20x objective. Images were captured as a large image with stitching. Image analysis was using the NIS Elements software.

### Fluorescent In Situ Hybridization

Whole brain was collected and flash frozen in liquid nitrogen. For sectioning, brains were embedded in OCT and frozen on cold isopentane and stored at -80°C. Tissue was sliced on a cryostat in 12 or 20 µm sections and stored at -80°C until processing. Fluorescent *in situ* hybridization was performed according to the manufacturer’s instructions using the RNAScope Fluorescent Multiplex Reagent Kit (Advanced Cell Diagnostics). Genes analyzed were *tyrosine hydroxylase* (ACDBio Cat#317621), *glutamic acid decarboxylase 2* (ACDBio Cat#439371), and *Slc30a10* (ACDBio Cat#484291). Coverslips were placed on each slide, sealed with DAPI-Fluoromount G (Southern Biotech, Cat# 0100-20), and allowed to dry overnight before imaging. Images were captured using a Nikon swept-field confocal with an inverted Nikon TiE microscope. The objective was a Nikon X20 or X60 oil-immersion objective. Images were captured as a large image with stitching.

### Surgical Procedures

All surgical procedures were performed using aseptic techniques. Surgeries were performed similarly to procedures previously described ^47,48^. Briefly, mice were anesthetized with isoflurane (2.5 - 3% for induction, 1 - 2% for maintenance) and secured on a stereotaxic apparatus (David Kopf Instruments, Tujunga, CA). A guide cannula (8 mm long, 20 gauge; Ziggy’s Tubes and Wires, Inc., Sparta, TN) was implanted to target the dorsal striatum using the following coordinates: +2.3 mm lateral to Bregma, +0.7 mm anterior to Bregma, 1.5 mm ventral to the skull surface. The guide cannula was secured using one stainless-steel screw (Antrin Minature Specialties, Inc., Fallbrook, CA) and dental cement. A wire loop for tethering during *in vivo* microdialysis was also secured to the skull with the dental cement. An obturator was inserted into the guide cannula to prevent the cannula from clogging. Mice were given a local analgesic (bupivacaine, 5 mg/kg; Covetrus North America) and a general analgesic (Rimadyl, 5 mg/kg body weight; Zoetis, Parsippany, NJ) subcutaneously during surgery. Following surgery, mice were allowed to recover on heat until fully awake and sternal. All mice were singly housed and provided with food and water ad-libitum. Mice were given general analgesic for 2 days post-surgery and as needed until microdialysis. General health and body weight were monitored daily.

### In Vivo Microdialysis

Microdialysis probes (3 mm active area) were constructed similar to Job et al. (2006) ^49^. Probes were inserted under light anesthesia (1 - 2% isoflurane) at least four days post-surgery. Following probe insertion, animals were placed in individual microdialysis chambers, and normal artificial cerebrospinal fluid (ACSF; 149 mM NaCl, 2.8 mM KCl, 1.2 mM CaCl_2_2H_2_0, 1.2 mM MgCl_2._6H_2_0, 5.4 mM D-glucose, and 0.05 mM ascorbic acid) was perfused at 0.2 µL/ min overnight. The following morning, the flow rate was increased to 1 µL/min at least 2 hours prior to sample collection. Sample collection occurred at least 12 hours after probe insertion. Three baseline samples were collected at 20-min intervals. For the high KCl sample, animals were perfused with ACSF containing 50 mM KCl (NaCl concentration was adjusted to maintain isotonicity) for 10 min, then returned to normal ACSF for the remaining 20 min. At least three additional baseline samples were collected. After the final baseline sample collection, calcium-free ASCF was perfused for at least 2 hours and samples collected to determine the calcium-dependency of dopamine release. All samples were frozen on dry ice immediately after collection and stored at -80°C until analysis by high-performance liquid chromatography.

### Liquid Chromatography

Concentrations of dopamine and DOPAC were determined using high-performance liquid chromatography (HPLC) with electrochemical detection. For dopamine and DOPAC analysis of striatal tissue, tissue preparation was performed similar to previous work ^50^. Briefly, the striatum was dissected by hand, flash frozen in liquid nitrogen, and stored at -80°C. To prepare tissue for HPLC analyses, 300 µL of 0.2 M perchloric acid per 10 mg of wet tissue was added. Samples were sonicated and kept on ice until injection. Samples not used for immediate injection were stored at -80°C. The mobile phase consisted of 0.0269 g 1-octanesulfonic acid sodium salt monohydrate (OSA), 3.0110 g citric acid monohydrate (Fluka), 0.02 g ethylenediaminetetraacetic acid disodium salt (EDTA), and 1.0170 g sodium dihydrogen phosphate dihydrate dissolved in 1 L MilliQ water, pH 4.6, and 7% (v/v) methanol. A Luna C18(2) LC Column (3 µm particle size, 100 Å, 150 × 2 mm; Phenomenex, Torrence, CA), guard column (BDS-Hypersil-C18, 5 µm 20 × 2.1 mm), and the VT03 (2 mm glassy carbon electrode) were used. The flow rate was set to 0.2 mL/min.

For dopamine analysis in dialysates, the mobile phase was dissolved in 1L MilliQ water, pH 5.6, and 12% methanol. A Luna C18(2) LC Column (3 µm particle size, 100 Å, 50 × 1 mm; Phenomenex, Torrence, CA) and SenCell (2 mm glassy carbon electrode, salt-bridge reference; Antec Scientific, The Netherlands) were used. The flow rate was set to 0.1 mL/min. Samples were thawed immediately before injection.

Chromeleon 6.8 Data System Software (Thermo Fisher Scientific, Waltham, MA, USA) was used for all data acquisition.

### ICP-MS Metal Analyses

ICP-MS metal analyses from bulk tissue samples were performed as described in our previous papers ^27,29,30^. Brain samples were either brain punches or a section of the midbrain.

### LA-ICP-MS

For LA-ICP-MS, brains were harvested at PND 21 from mice that had received daily oral Mn treatment from birth as described above, and 20 µm thick coronal sections on glass slides were prepared. Sections corresponding to the striatum/globus pallidus and substantia nigra were processed for LA-ICP-MS. These sections were identified by an atlas, and RNAscope imaging was performed on alternate serial sections to validate presence of GABAergic or dopaminergic neurons. LA-ICP-MS analyses were performed using a Photon Machines Analyte Excite laser ablation system coupled to a Thermo Element XR magnetic-sector ICP-MS, with analysis parameters of 20 µm laser spot size, 20 Hz pulse rate, and 40 µm/sec laser raster rate, and raster lines of ∼4000 µm and six lines/section. The laser was rastered across each hemisphere in a dorsoventral direction for the striatum/globus pallidus samples, and across both hemispheres in a lateral direction for the substantia nigra samples.

### Chemicals

Unless otherwise noted, all chemicals were from Thermo Fisher Scientific or Sigma-Aldrich.

### Statistical Analyses

All statistical analyses were performed using the PRISM 9 software (Graphpad Inc., La Jolla, CA). Comparisons between multiple groups were performed using a one- or two-way ANOVA with appropriate *post-hoc* analyses. Comparisons between two groups were performed using a Student’s *t* test. Comparisons that involved multiple groups and time points were performed using a two-ANOVA with repeated measures and appropriate post-hoc analyses. For all testing, *p* < 0.05 was considered statistically significant. *Asterisks* in figures indicate a statistically significant difference.

## Acknowledgements

Supported by NIH/NIEHS R01 ES024812 (S.M.) and NIH/NINDS F99 NA124142 (C.A.T.).

## Notes

### Competing Interest Statement

The authors have declared no competing interest.

