## Supplemental Fig 1 for "Activity of the manganese efflux transporter SLC30A10 in dopaminergic but not GABAergic neurons protects against neurotoxicity"

**Supplemental Materials for Taylor *et al***

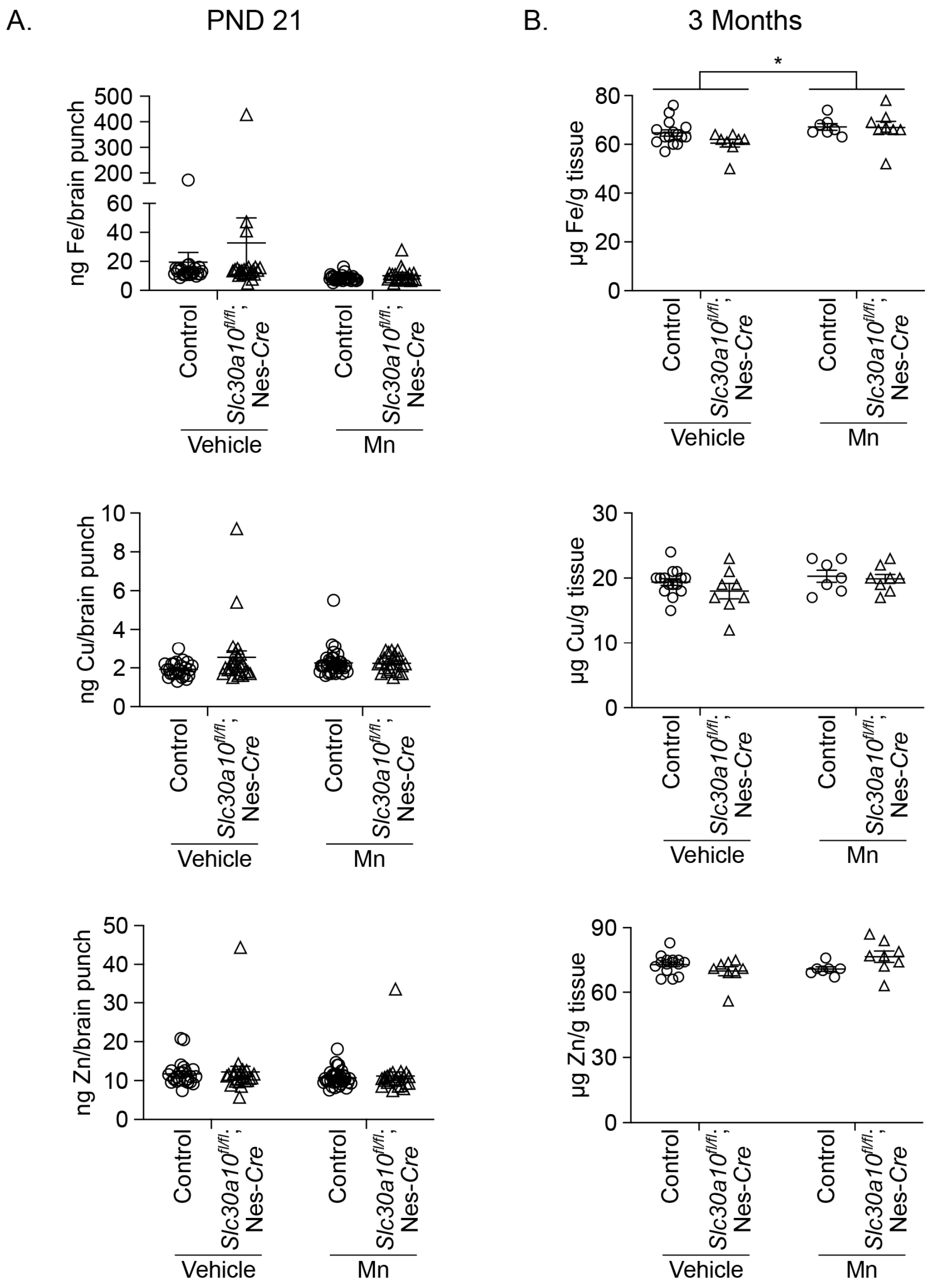


**Figure S1.** Levels of other metals (Fe, Cu, and Zn) were in vehicle- or Mn-treated pan-neuronal/glial *Slc30a10* knockouts were comparable to littermate controls.

***A.*** ICP-MS to measure levels of Fe, Cu, and Zn in brain punches of animals from **Fig.1B**. Mean ± SE.

***B.*** ICP-MS to measure levels of Fe, Cu, and Zn in brain tissue of animals from **Fig.1D.** Mean ± SE. *, $p < 0.05$ by two-way ANOVA for a main effect of Mn treatment.
